# Bioinformatics analysis of the potential biomarkers of Multiple Sclerosis and Guillain-Barré syndrome

**DOI:** 10.1101/2024.05.29.595759

**Authors:** Tyler Kwok, Tajah Huerta-White, Karl Briegel, Aaisha Singh, Suneetha Yeguvapalli, Kumaraswamy Naidu Chitrala

## Abstract

Recent research emphasizes the intricate interplay of genetics and epigenetics in neurological disorders, notably Multiple Sclerosis (MS) and Guillain-Barre Syndrome (GBS), both of which exhibit cardiovascular dysregulation, with GBS often featuring serious bradyarrhythmias requiring prompt recognition and treatment. While cardiovascular autonomic dysfunction in MS is typically less severe, orthostatic intolerance affects around half of MS patients. Their distinction lies in their autoimmune responses, MS is an autoimmune disease affecting the central nervous system, causes demyelination and axon damage, leading to cognitive, ocular, and musculoskeletal dysfunction. In contrast, GBS primarily affects the peripheral nervous system, resulting in paralysis and respiratory complications. Despite their differences, both diseases share environmental risk factors such as viral infections and Vitamin D deficiency. This study aims to explore shared gene expression pathways, functional annotations, and molecular pathways between MS and GBS to enhance diagnostics, pathogenesis understanding, and treatment strategies through molecular analysis techniques. Through the gene expression analysis, five significant genes were found UTS2, TNFSF10, GBP1, VCAN, FOS. Results shows that Common DEGs are linked to apoptosis, bacterial infections, and atherosclerosis. Molecular docking analysis suggests Aflatoxin B1 as a potential therapeutic compound due to its high binding affinity with common differentially expressed proteins.

## Introduction

In recent years, neurological disorders have been a steady topic at the forefront of research studies due to the intricate interactions between genetics and epigenetics in the pathogenesis of neurological disorders. Among these disorders, Multiple Sclerosis (MS) and Guillain-Barré Syndrome (GBS) present unique challenges and unknowns for clinicians and researchers alike. Studies have shown that both multiple sclerosis and Guillain-Barré syndrome encounters cardiovascular dysregulation. [1] In GBS, cardiovascular dysregulation, including serious bradyarrhythmias, is common and may require early recognition and appropriate therapy. Cardiovascular autonomic dysfunction in MS is typically of minor clinical importance, but orthostatic intolerance may affect around 50% of patients. Both diseases present distinct clinical properties and have an effect on the cardiovascular autonomic dysfunction, making them compelling subjects for comparative analysis at the molecular level.

Multiple Sclerosis is a non-traumatic neurological autoimmune disease (NAD) that primarily affects young adults [2]. The disabling disease is characterized by the damage and demyelination of Oligodendrocytes as well as progressive axon damage, by leukocytes. These processes often present as symptoms such as progressive cognitive disfunction, ocular impairment, and musculoskeletal dysfunction, among others. For Guillain-Barré syndrome (GBS), a rare NAD, leukocytes target the myelin sheaths and axons of Schwann Cells [3]. In the most severe cases, GBS may lead to partial or complete paralysis, affecting breathing. The distinction between MS and GBS results from the underlying pathology of their autoimmune responses: MS primarily affects the central nervous system, while GBS predominantly triggers a response in the peripheral nervous system [2, 3]. While the diseases may exhibit similarities, they differ significantly in terms of conditions and treatment approaches. MS typically necessitates lifelong management of symptoms, progression, and relapse, while GBS is commonly acute and may require post therapy rehabilitation to regain lost function [2, 3]. Early onset treatment of GBS may also lead to more successful symptom management, reducing its severity and duration of treatment. The underlying causes of the diseases are not completely understood. However, there is strong evidence suggesting that epigenetics may exert a greater influence on susceptibility than an individual’s genetics [4]. Studies have identified common environmental factors including viral infection, Vitamin D/UV-B deficiency and geographical location, and smoking. Particularly, Epstein-Barr Virus and Vitamin D deficiency have been highly linked to susceptibility between diseases [2, 5, 6].

This study looks to analyze the complex expression pathways of both diseases to determine whether they share differentially expressed genes, functional annotations, or molecular pathways. Identifying novel biomarkers and pathways may provide insight into the intricate mechanisms behind the diseases to abate gaps in diagnostics, pathogenesis, management, and treatment. Utilizing various bioinformatic tools and advanced molecular analysis methods, we will compare genome-wide gene expression profiled in patient datasets for MS and GBS. This research seeks to unveil the molecular markers driving the initiation, progression, and modulation of these neurological diseases.

## Results

Identification of DEGs and obtaining common DEGs

Figure 1 shows the flow chart of the overall methodology that were used for this study. In the GSE235357 dataset, a comprehensive analysis revealed the identification of 365 DEGs, among them 209 genes were up-regulated and 156 are down regulated (Figure 2a). In the GSE31014 dataset, 148 DEGs were identified, 52 genes are up-regulated and 96 genes are down-regulated (Figure 2b). We identified 5 common DEGs: UTS2,TNFSF10,GBP1,VCAN and FOS as shown in the Venn diagram (Figure 3).

**Figure 1.**
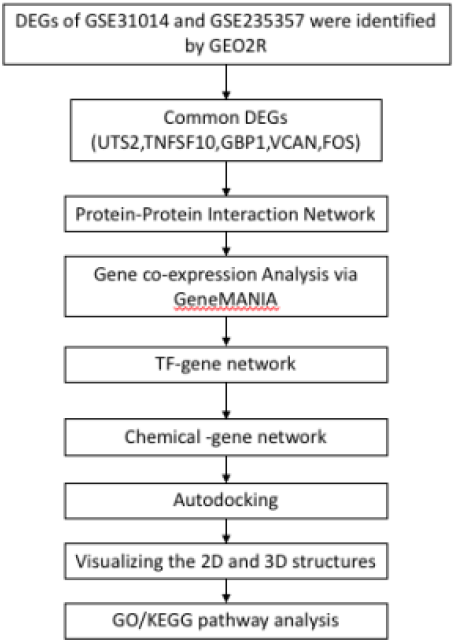
Flowchart of the bioinformatic analysis conducted for MS and GBS

**Figure 2.**
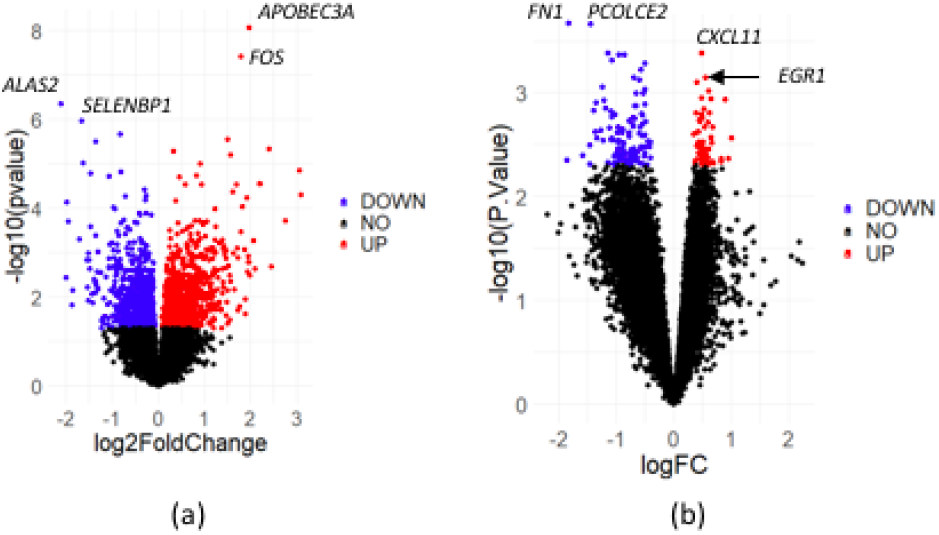
Volcano diagrams of Differential Expression Results for MS(a) and GBS(b) Datasets.

**Figure 3.**
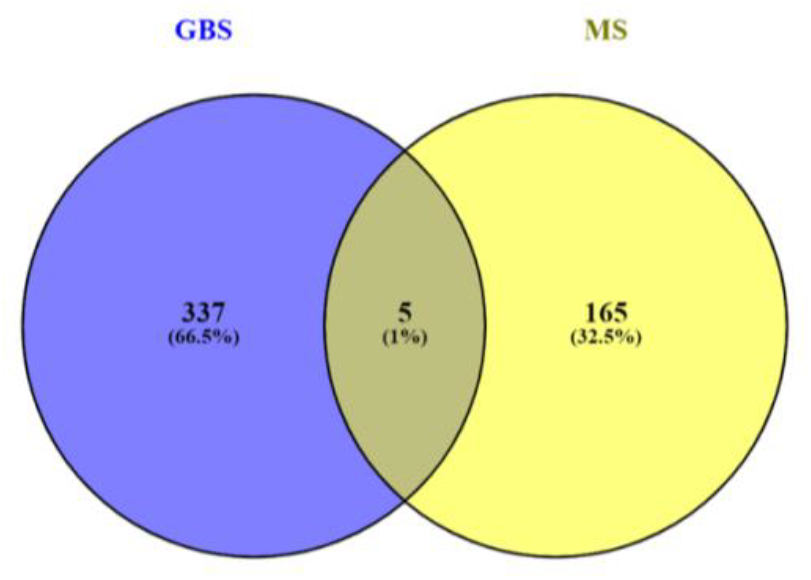
Venn Diagram of Shared Significant Differentially Expressed Genes.

**Figure 4.**
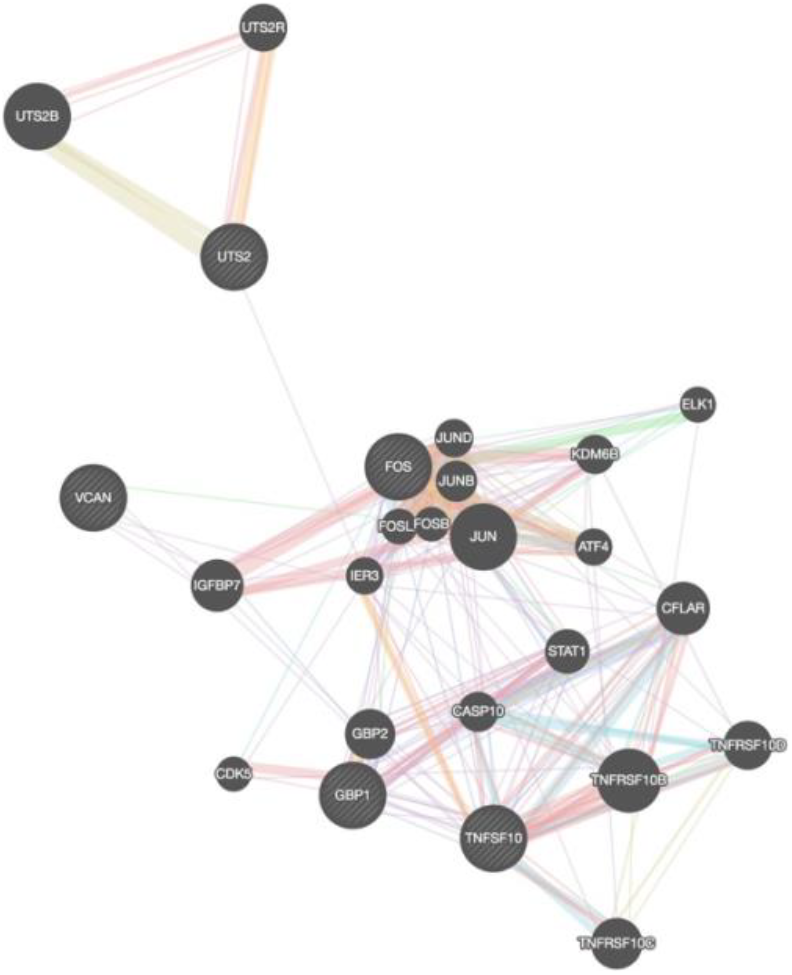
Protein-protein interaction analysis of the five common DEGs via Gene Mania. Red Colored line resembles physical interaction, purple colored line resembles co-expression, orange colored line resembles predicted, blue colored line resembles co-localization, green colored resemble genetic interaction and light blue resembles pathways

### Construction of PPI network

A The PPI network of the common DEGs was constructed using STRING database (supplementary figure), containing 5 nodes and 1 edges. The interaction shows that GBP1 and TNFSF10 are coexpresesed and they are enriched in Sleep regulation and Photodynamic therapy-induced AP-1 survival signaling pathways. We used the GeneMANIA to get gain further detailes on the gene interactions of the five DEGs. The results show that there is a 77.64% and 8.01% of genes are Co-expression, 17.39% are Shared protein domains (Figure 5). In supplementary figure —, we identified the signaling relationship of the 5 common DEGs. The results shows that FOS form complex bind with AP1 (score: 0.951), and (-)-anisomycin upregualtes and lead to chemical activiation of FOS (score: 0.8). The upregualtion of MAPK3 (score: 0.707) and RPS6KA1(score: 0.55) lead phosphrlyation of FOS. ETS1(score: 0.706) and CREB1(score: 0.652) up-regulates quantity by expression lead to transcriptional regulation of FOS.

**Figure 5.**
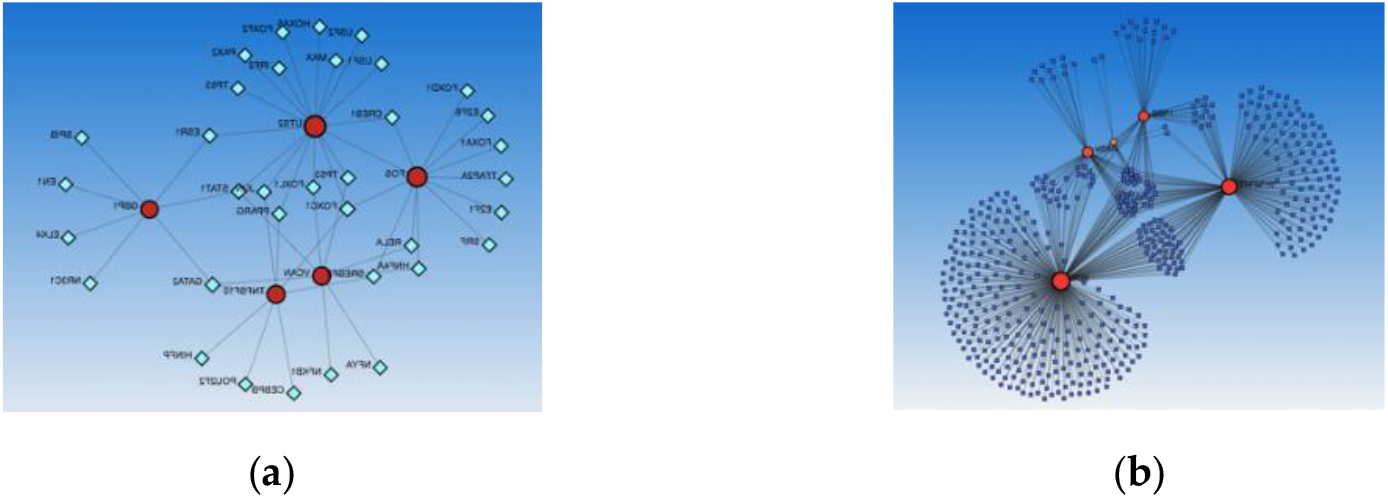
Constructed a network analysis for Target transcription factor(a) and Chemical(b) to the 5 common DEGs respectively. The red nodes repre-sent the 5 common DEGs, the light blue represent the target transcription factor and the dark blue represent the chemical compounds.

**Figure 6.**
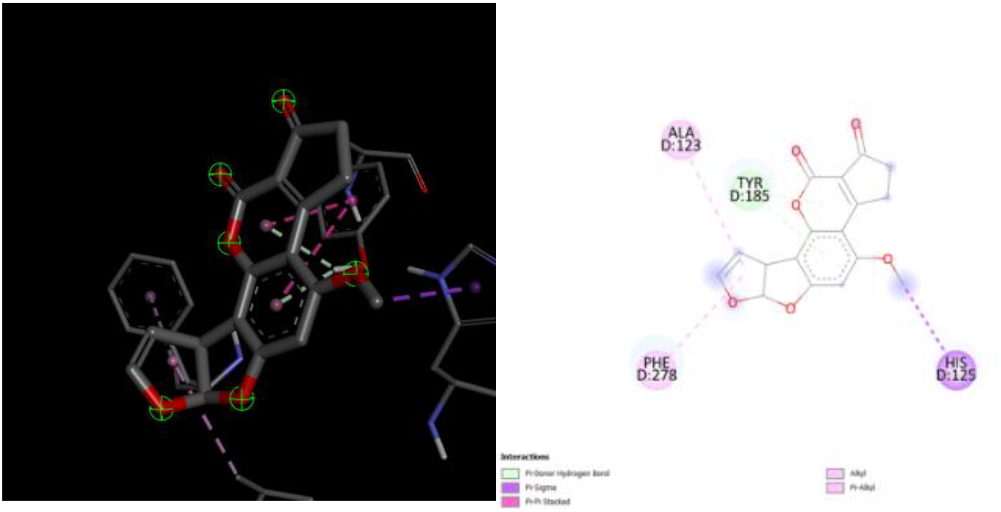
2D and 3D structure of Aflatoxin B1 binding to PDB structure: 61kz (GBP1)

A network analysis is performed to identify the targeted transcription factor to DEGs and chemical to DEGs interaction. The network for targeted transcription factor to DEGs included 40 Nodes and 51 Edges. FOXC1, GATA2 and STAT1 has the highest degree(3) of interaction with the DEGs. For our chemical to DEGs interaction, it has 499 Nodes and 665 Edges, Aflatoxin B1, (+)-JQ1 compound, Silicon Dioxide and Tetrachlorodibenzodioxin has the highest degree (5) of interaction.

### Binding affinity of chemical and cDEG interactions

We selected Aflatoxin B1, (+)-JQ1 compound and Tetrachlorodibenzodioxin from the chemical-DEGs network analysis to examine the binding affinity with gene GBP1(PDB structure: 6k1z), TNFSF10 (PDB structure: 5CIR) and FOS (PDB structure: 1S9K). The results shows that Aflatoxin B1(-8.2) has the highest binding affinity with GBP1 following with (+)-JQ1 compound(-7.4) and Tetrachlorodibenzodioxin(-7.3). As shown in Figure 7a, Aflatoxin B1 showed one hydrogen bonds with LYS520 residues, two pi-pi T shaped interaction at TYR52, one pi-pi T shaped interaction at HIS527, one alkyl at ALA385 and four pi-alkyl interaction at TYR524 and HIS527. (+)-JQ1 compound (Figure 7b) formed one Pi-anion interaction GLU389, one pi-sigma interaction with ALA385 and two pi-Alkyl interaction at TYR524 and UNK1. Tetrachlorodibenzodioxin formed three Pi-alkyl interactions with HIS378 and two at ALA385. There are six alkyl, two at ALA385, two at LEU531, one at LEU528 and CYS589 respectively. Three carbon hydrogen bond is found at HIS378, LEU528 and TYR524, three pi-pi stack interaction is formed at TYR524, one Pi-pi T shaped interaction is form with HIS527.

**Figure 7.**
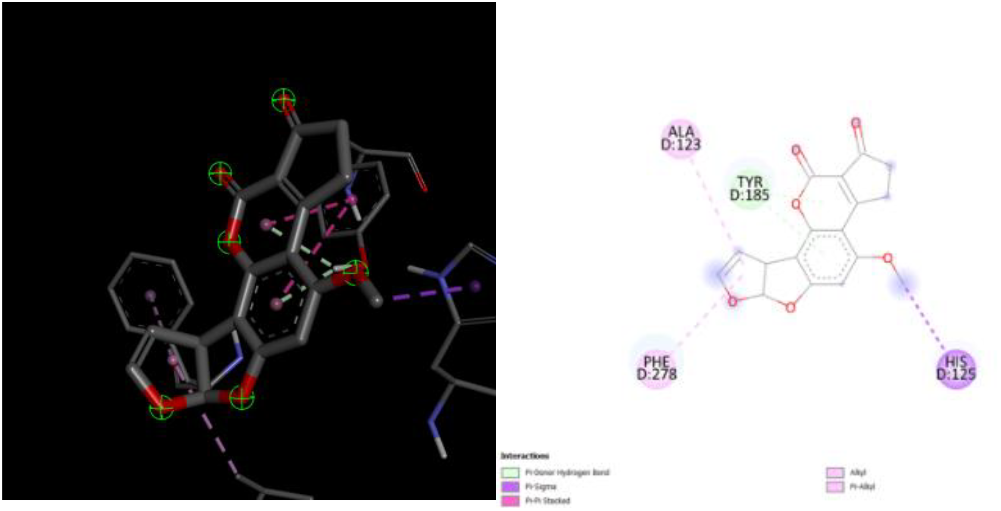
2D and 3D structure of Aflatoxin B1 binding to PDB structure: 5CIR (TNFSF10)

**Figure 8.**
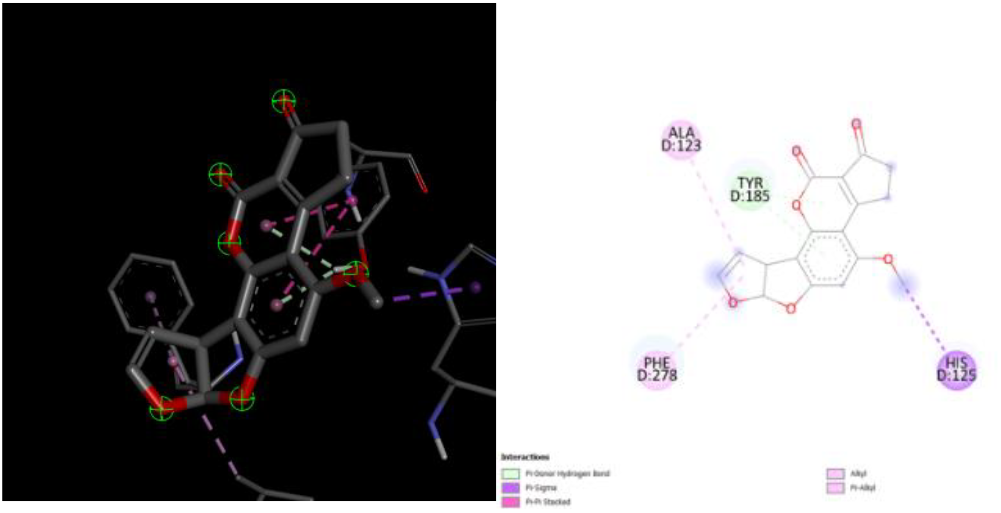
2D and 3D structure of Aflatoxin B1 binding to PDB structure: 1s9k (FOS)

**Figure 9.**
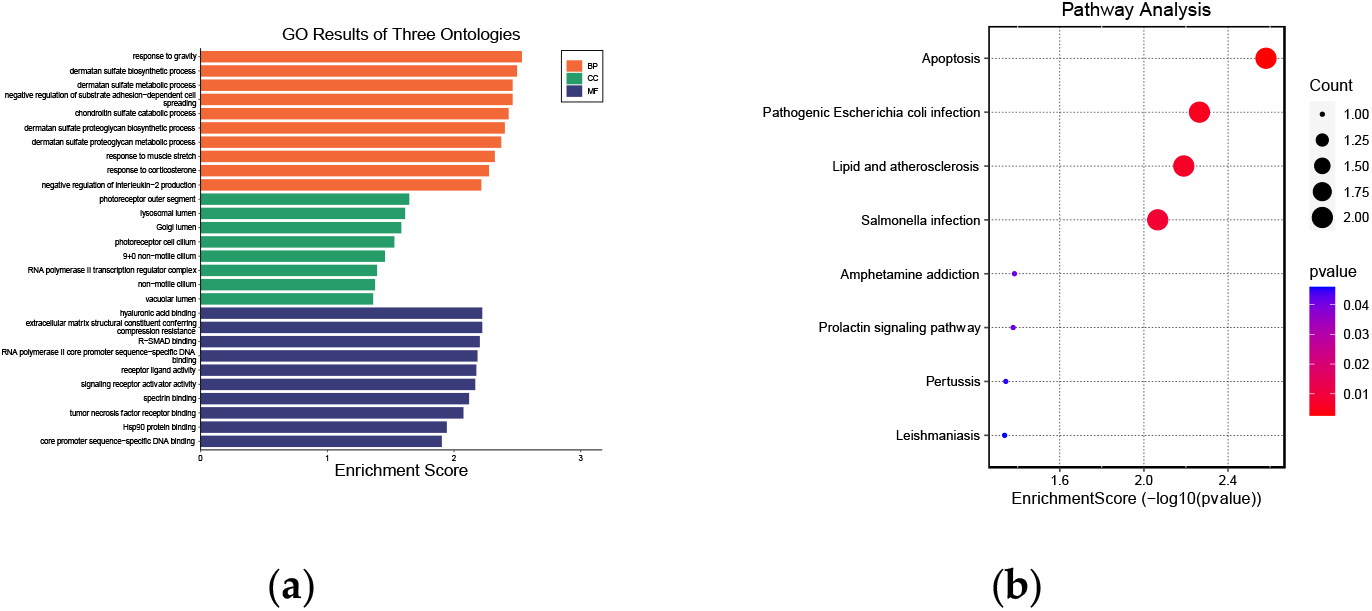
Functional enrichment analysis for the 5 common DEGs. A) is the gene ontology analysis b) is the KEGG pathway analysis.

TNFSF10 has the highest binding affinty with Aflatoxin B1 (-6.9), following with (+)-JQ1 compound(-6.8) and Tetrachlorodibenzodioxin (-5.7). Aflatoxin B1 has two Pi-Donor hydrogen bond at TYR185, one Pi-sigma interaction at HIS125, two Pi-pi stacked at TYR185, one alkyl at ALA123 and one pi-Alkyl interaction at PHE278. (+)-JQ1 compound. (+)-JQ1 compound showed two hydrogen bond at GLN205,ASN262, one Carbon Hydrogen bond at LYS204 and one Pi-anion interaction at ASP203. One alkyl and pi-alkyl interaction are found at LYS204. Tetrachlorodibenzodioxin has one carbon Hydrogen bond and one Pi-donor hydrogen bond at HIS125 and TYR185 respectively. Two Pi-Pi stcked interaction with TYR185 and two alkyl at ALA123.There is 8 pi-alkyl interaction at HIS125,PHE163,TYR243,PHE274,PHE278,ALA123 and two at TYR185.

FOS has the highest binding affinty with Aflatoxin B1 (-8.1) following with (+)-JQ1 compound (-8.1), and Tetrachlorodibenzodioxin (-7.0). Aflatoxin B1 has one hydrogen bond at TYR474, one pi-anion hydrogen bond at DT5017 and three pi donor Hydrogen bond: one at TYR424 and two at SER570. One alkyl at ARG572 and two pi-alkyl interaction at DT5016 and DT5017. (+)-JQ1 compound has one hydrogen bond at GLN404, three alkyl at LEU187,ARG556 and VAL558. There is four pi-alkyl interaction: two at ALA184, one at VAL558 and one at LYS188. Tetrachlorodibenzodioxin has three Pi-donor Hydrogen bond at SER570, one at TYR424 and one at GLN51. Two pi-anion interaction at DT5017, one pi-sigma interaction at SER570 and one pi-pi T-shaped at TYR424. One pi-alykl interaction at DT5016, one at TYR474 and two at TYR424. It also has five alkyl at ARG537,ARG421,LYS520 and CYS569.

### SwissTargetPrediction

Results from molecular target prediction by SwissTargetPrediction tool provided several possible interacting targets for our chemical compounds. Aflatoxin B1 was predicted to interact with Kinase (73.3%), Lyase (6.7%) ,other cytosolic protein (6.7%), family A G protein coupled receptor (6.7%) and Oxidoreductase (6.7%). The results from the target receptors only show low probabilities of interaction. (+)-JQ1 compound was predicted to interact with Kinase (26.7%), Reader (26.7%), family A G protein coupled receptor (13.3%), electrochemical transporter (6.7%), enzyme (6.7%), Family C G protein coupled receptor(6.7%), Cytochrome P450(6.7%) and Protease (6.7%). (Supplementary Fig. 3). It has a high probability of interaction with Bromodomain-containing protein 4, Bromodomain-containing protein 2, Bromodomain testis-specific protein and Bromodomain-containing protein 3. Tetrachlorodibenzodioxin was predicted to interact with protease(20%), enzyme(20%),other cytosolic protein (20%), oxidoreductase (13.3%), kinase (13.3%), electrochemical receptor (6.7%) and electrochemical receptor. It has a high probability of interaction with Aryl hydrocarbon receptor and Vascular endothelial growth factor receptor 1.

### Functional enrichment analysis

Functional enrichment analysis was performed for the 5 common DEGs. The KEGG pathway analysis shows that the 5 DEGs were significantly enriched in Apoptosis, pathogenic Escherichia coli infection, Lipid and atherosclerosis and Salmonella infection. The Gene Ontology shows that Biological Process enriched in response to gravity, dermatan sulfate biosynthetic process and dermatan sulfate metabolic process. Cellular composition is enriched in Photoreceptor outer segment, lysosomal lumen and Golgi lumen. Molecular function is enriched in hyaluronic acid binding, extracellular matrix structural constituent conferring compression resistance and R-SMAD binding.

## Discussion

Multiple sclerosis (MS) is a complex autoimmune disease which in the central nervous system causes tissue inflammation, demyelination, and neurodegeneration of nerve fibre [7]. Guillain-Barré Syndrome (GBS) is an uncommon neurological condition in which the body gets confused and its immune system starts mistakenly targeting part of its peripheral nervous system, which is the network of nerves lying outside the brain and the spinal cord. One of the key aspects of MS and GBS bioinformatic research is the ability to identify biomarkers (markers which help to diagnose, define prognosis and comprehend the pathogenesis) and find therapeutic targets. We performed DEG analysis for the healthy and MS Baseline samples; and samples from a normal control and a GBS patient. By comparing both datasets, we found five significant DEGs that are commonly expressed (UTS2, TNFSF10, GBP1, VCAN, FOS)

UTS2(Urotensin 2) is a vasoactive peptide, involved in stimulating the inflam-matory process[8] The immune system is the major player in GBS in-flammation because it is turning against the myelin sheath of the nerve. The evidence of an increased level of UTS2 might indicate that GBS is an inflammatory condition and is thus a potential marker of disease activity or severity. Additionally, UTS2 is re-sponsible for brain inflammation and it may play a role in multiple sclerosis (MS) causing CNS inflammation.

The 10th group of the TNF superfamily is a protein called TNFSF10, which is also commonly known as TRAIL. The molecules have their influence on the maturation of the apoptotic signals as well as inflammatory pathways [9] TNF-superfamily member 10 (TNFRSF10), which is a disorder that has a connection with autoimmune disease, is high in the lesions of multiple sclerosis patients. To date, an extensive body of research has shown that this superfamily plays a role in the pathogenesis of MS, regulating disease progression by modulating both the peripheral immune response and interactions between CNS-resident cells and effector immune cells within the CNS[10]. This can be employed to elucidate the contributions of the members to their pathogenesis, as well as a predictor for disease progression. Moreover TNFSF10 is expressed by activated immune cells and triggers apoptosis (programmed cell death) which is used by the immune system to regulate the response to infection.[9] In GBS, the consequence of aberrant apoptosis may be a reduction level of nerve cell death or modulation of immune cells.

GBP1 (Guanylate Binding Protein 1) is a protein that belongs to a family of vi-rus-induced GTPases which regulate the immune system and prevent unnecessary or exaggerated immune reactions. The brain is a tissue of MS patients characterized by inflammation, it is the source of inflammation in the brain. They have found this out by high levels of GBP1 in the serum of MS patients in which case they now can use GBP1 as a biomarker to follow the inflammation status and therapy response of MS patients [11]. Additionally, GBP1 is an antiviral protein whose role is not only limited to immunomodulation [12]. Hence, its overexpression might be an asset of a hyper-reactive or engaged immune system that is trying to pro-tect the body from an imaginary pathogen invasion, often starting the entire episode. Thus increased GBP1 can be an indicator of activation of the immune system in GBS in terms of the pathophysiological processes.

Versican is a large extracellular matrix proteoglycan, and it takes part in cell ad-hesion, proliferation, as well as migration [13]. Versican has been thought to be involved in inflammation in several studies. The VCAN protein possibly could be involved in forming the extracellular matrix of nerves and nearby tissues, and this might have a significant impact on nerve function and repair. VCAN protein lev-els may give some information about the maximum degree of nerve injury and the possibility of restoring the nerves. They can also be used as markers of disease severity or recovery in GBS and MS.

FOS is a proto-oncogene, a gene which is associated with cell division and matu-ration and those processes which are related to cancer. The link between the shift of FOS degree and inflammatory reactions of multiple sclerosis has been already proven [14]. In MS lesions, FOS expression has been well detected as a potential biomarker to track the disease conditions and dynamical progression. Moreover, FOS belongs to the AP-1 transcription factor complex which is responsible for the regula-tion of different cellular processes – proliferation, differentiation, and survival [15]. It is rapidly induced by various signals, including cytokines and growth factors that are abundant when inflammation is present. In GBS, FOS can be involved in the regulation of genes which would determine response to infections or nervous system repair, thereby acting as a sign of disease activity or response to treatment. The KEGG pathway enrichment results showed that common DEGs were mainly associated with Apoptosis, Pathogenic Escherichia coli infection, Salmonella infection and lipid and atherosclerosis. These results also provided significant clues to studying molecular interactions in the progression of MS and GBS.

The relationship between chemical compounds and proteins is essential for com-prehending how biological processes are regulated and for advancing the development of therapeutic drugs[16]. The results from our molecular docking analy-sis, we found that Aflatoxin B1 has the highest binding affinity among GBP1, FOS and TNFRSF10 compared to the other chemical compound. Additionally, Aflatoxin B1 are soluble in water and high level of absorption level compared to other chemical compounds (supplementary figure 7). It suggest that Aflatoxin B1 could be the most effec-tive chemical compound when developing drugs for treating MS and GBS. More stud-ies are needed to validate the effect of the chemical compounds and refined as drugs.

## Conclusion

In conclusion, UTS2, TNFSF10, GBP1, VCAN, FOS play a crucial role in the pathogenesis and clinical association of MS and GBS. These genes could be used as novel diagnostic and prognostic biomarkers. Aflatoxin B1 was found be an essential chemical com-pound for developing therapeutic drugs when treating with MS and GBS.

## Materials and Methods

### Dataset preparation

In this study, we used bioinformatics methodology to find crucial differentially expressed genes (DEGs) and explain the potential function of the genes. Gene expression datasets related to Multiple Sclerosis and Guillain-Barré syndrome were obtained from the National Center for Biotechnolo-gy Information’s Gene Expression Omnibus (GEO) database (https://www.ncbi.nlm.nih.gov/geo/)[17] (accessed April 8th). We retrieved 10 Healthy donors and 10 Multiple Sclerosis samples from the GSE235357 (n=20), and 7 healthy controls samples and 7 GBS Guillain-Barré syndrome samples were retrieved from the GSE31014 dataset (n=14). Differential gene expression (DEG) analysis was performed be-tween(1) Healthy donor versus MS Baseline and (2) peripheral leukocytes from a normal control versus peripheral leukocytes from a GBS patient. A Volcano plot was created using ggplot2,ggrepel,dplyr and readrR packages

### Identification of DEGs and obtaining common DEGs

To identify the significant and overlapped DEGs between the GSE235357 and GSE31014 datasets, we used GEO2R (www.ncbi.nlm.nih.gov/geo/geo2r/)[18](accessed April 9th) with the following cut-off threshold values: log2(fold change [FC]) value>0 and p-value<0.005. Then we conducted Venn diagram analysis using VennDiagram R language package.

### Functional enrichment analysis

Gene Ontology and KEGG pathway enrichment analyses of the 5 common differential expressed genes were performed using the Data-base for Annotation, Visualization, and Integrated Discovery (DAVID; v6.8, http://david.ncifcrf.gov) (accessed April 10th)[19] database. SR plot (https://www.bioinformatics.com.cn/srplot) (accessed April 11th)[20] was used to generate graphs for Gene Ontology and KEGG pathway enrichment analyses for the 5 common significant genes with ggplot2 function in the R package.

### Construction of PPI network

We utilized STRING online analysis tool (http://www.string-db.org/)(accessed April 12th)[21] to analyze the protein-protein interactions of our common DEGs. Then, we use GeneMANIA (https://genemania.org/)[22](accessed April 13th) to validate and predict functions for our 5 common DEGs. For GeneMANIA, we focused on Homo sapiens as the target organism and selected interactions with a score greater than 0.4. Additionally, to explore signaling relationships associated the 5 DEGs, we utilized Signor 3.0 [23] (accessed April 14th) (https://signor.uniroma2.it/APIs.php)

### Regulatory networks

We utilized the NetworkAnalyst (https://www.networkanalyst.ca/)[24](accessed April 15th) online tool to visualize an interaction network between DEGs and transcription factors (gene-TF). This gene-TF network was built using data from JASPAR, a transcriptional factor binding site profile database. A chemical to DEGs interaction network was also generated with the use of Comparative Toxicogenomics Database.

### Molecular binding analysis

RCSB Protein Data Bank (PDB) was used to select the 3D structures of the 5 common DEGs. Discovery Studio Visualizer 3.0 (available at https://discover.3ds.com/discovery-studio-visualizer-download) (accessed April 17th) was utilized to prepare the protein for docking. This involved the removal of water and heteroatom compounds, as well as the addition of polar hydrogens. PubChem [25](https://pubchem.ncbi.nlm.nih.gov/) was used to download the structure of the chemical compounds Aflatoxin B1(Pubchem ID:186907), (+)-JQ1 compound(Pubchem ID: 46907787) and Tetrachlorodibenzodioxin(Pubchem ID:15625). The protein structures are input as macromolecules and transcription factor are input as ligands to identify their binding affinity using Virtual Screening software interface PyRx [26](https://pyrx.sourceforge.io/). The interaction will be visualized through Discovery Studio Visualizer 3.0. Then we analyze the druglikeness of the chemcial compound using SwissADME [27](http://www.swissadme.ch/index.php).

### Predicting Chemical targets

SwissTargetPrediction server (http://www.swisstargetprediction.ch/)[28](accessed April 16th)which combines both 2D and 3D similarity measures with our selected chemical compounds for knowledge-based prediction of potential targets.

## Acknowledgments Funding

National Institutes of Health grant R15NS116742 (AJF, VKN, SHG, WLS)

National Science Foundation grant R01AB123456 (AJF, SHG, WLS)

Arkansas Biosciences Institute (SHG, WLS)

## Author contributions

Conceptualization: AJF, JSS, WLS

Experimental recordings: AJF, VKN, SHG, WLS

Data analysis: AJF, JSS, WLS

Computational modeling: JSS

Figure creation: AJF, JSS, WLS

Writing: WLS, AJF, JSS

## Data and materials availability

Spike data and analysis code are publicly available without restriction on Figshare at doi: *TBD upon paper acceptance*. All data are available in the main text or the supplementary materials.

## Author contributions

Conceptualization: KNC

Experimental recordings: KNC, SY ,TH, AS, TK

Data analysis: KNC, SY ,TH, AS, TK

Computational modeling: KNC, SY ,TH, AS, TK

Figure creation: KNC, SY ,TH, AS, TK, KB

Writing: KNC, SY ,TH, AS, TK,KB

## Supplementary Materials

Supplementary Materials include seven Supplementary Figures

**Supplementary Figure 1.**
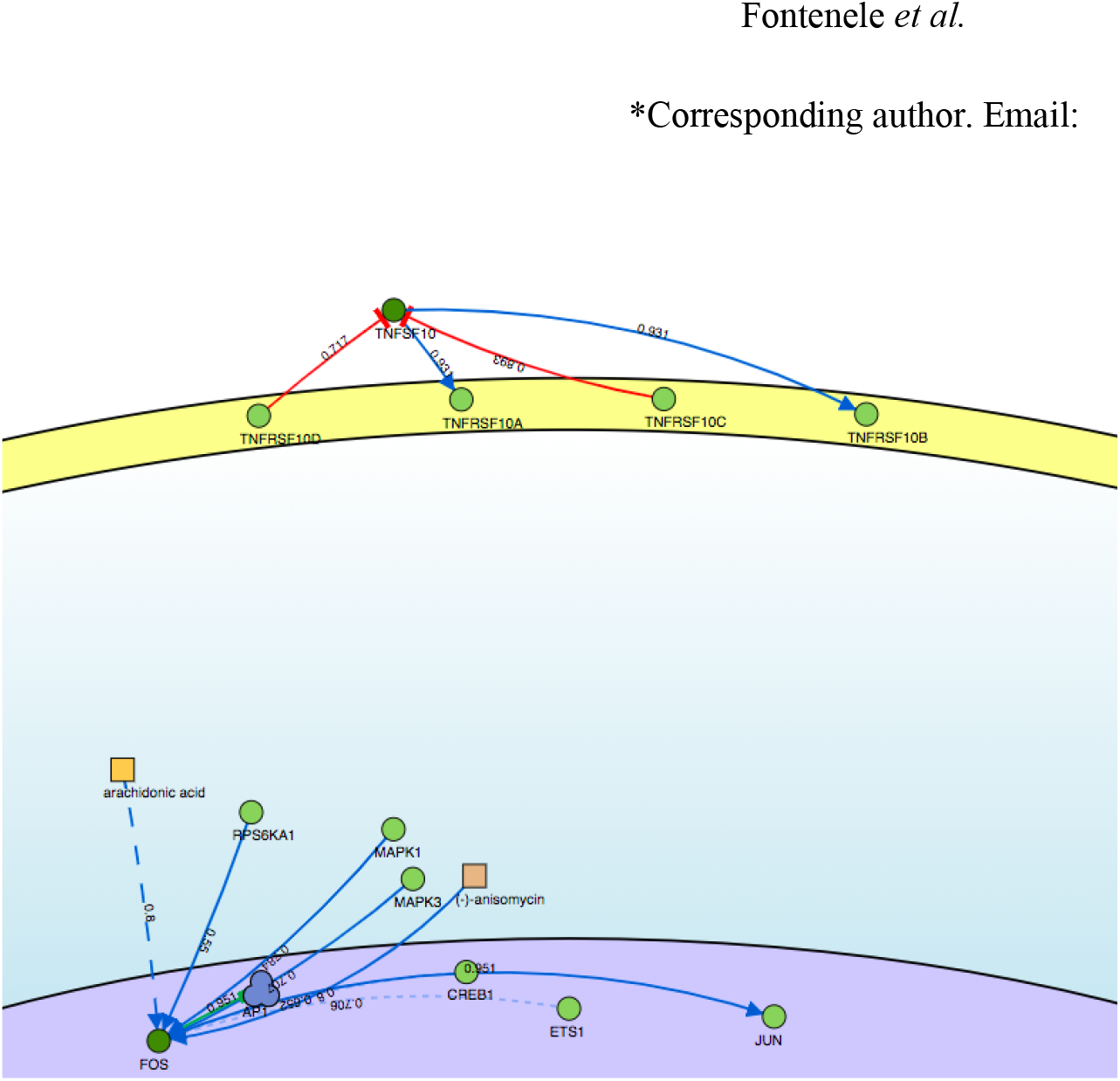
Analysis of Signaling pathways for the genes. Network showing the signaling pathways of the genes *FOS, TNFSF10*, and *VCAN* with a confidence score of 0.5 and above. Inhibited pathways are colored in red and activated pathways are colored in blue.

**Supplementary Figure 2.**
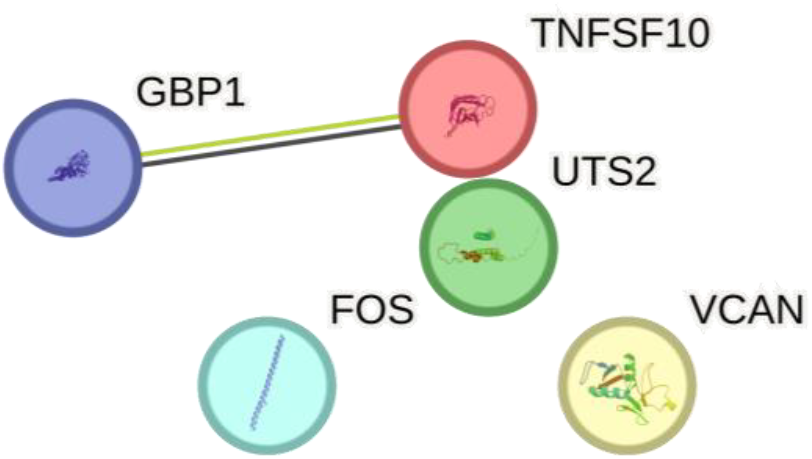
PPI network of genes the five common DEGS

**Supplementary Figure 3.**
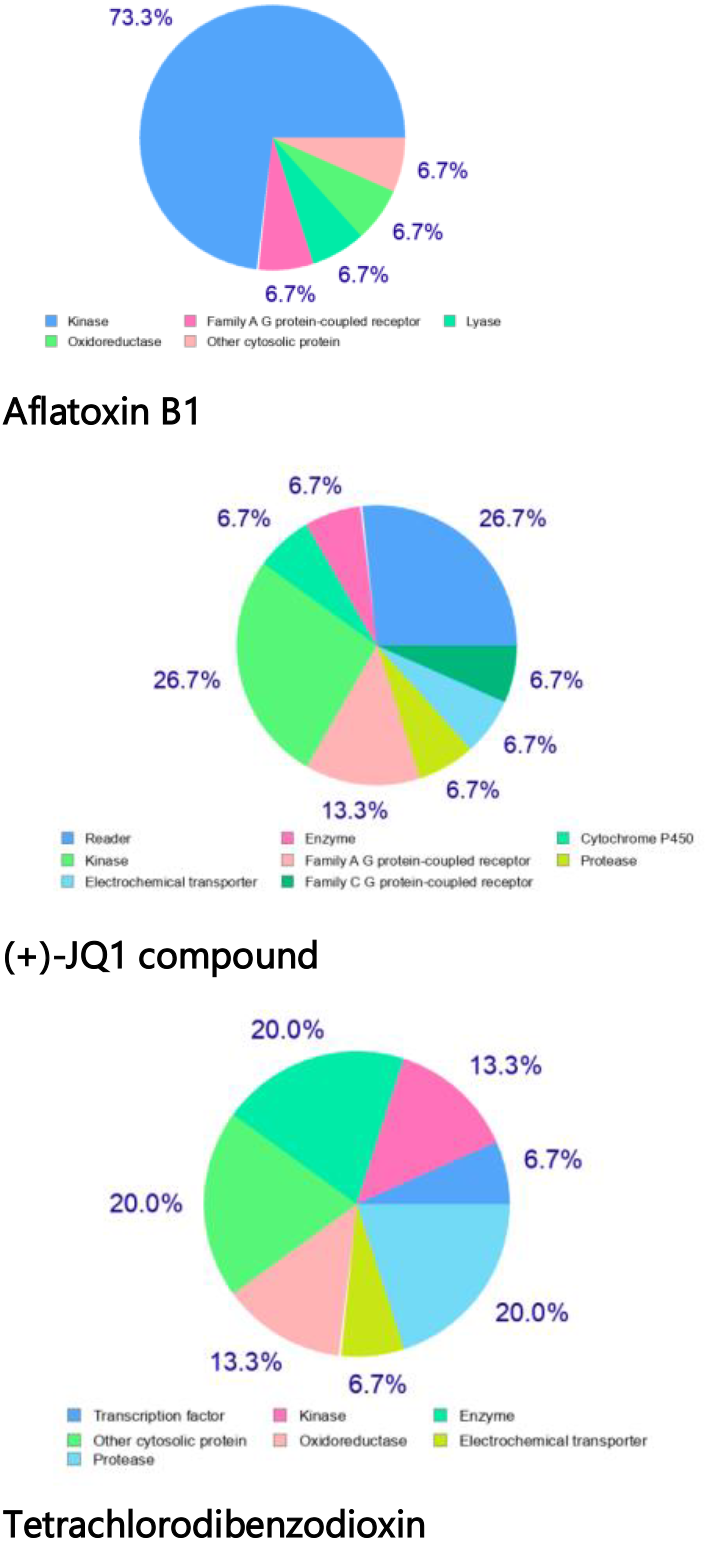
Pie chart of the potential target receptors for our selected chemical compounds

**Supplementary Figure 4.**
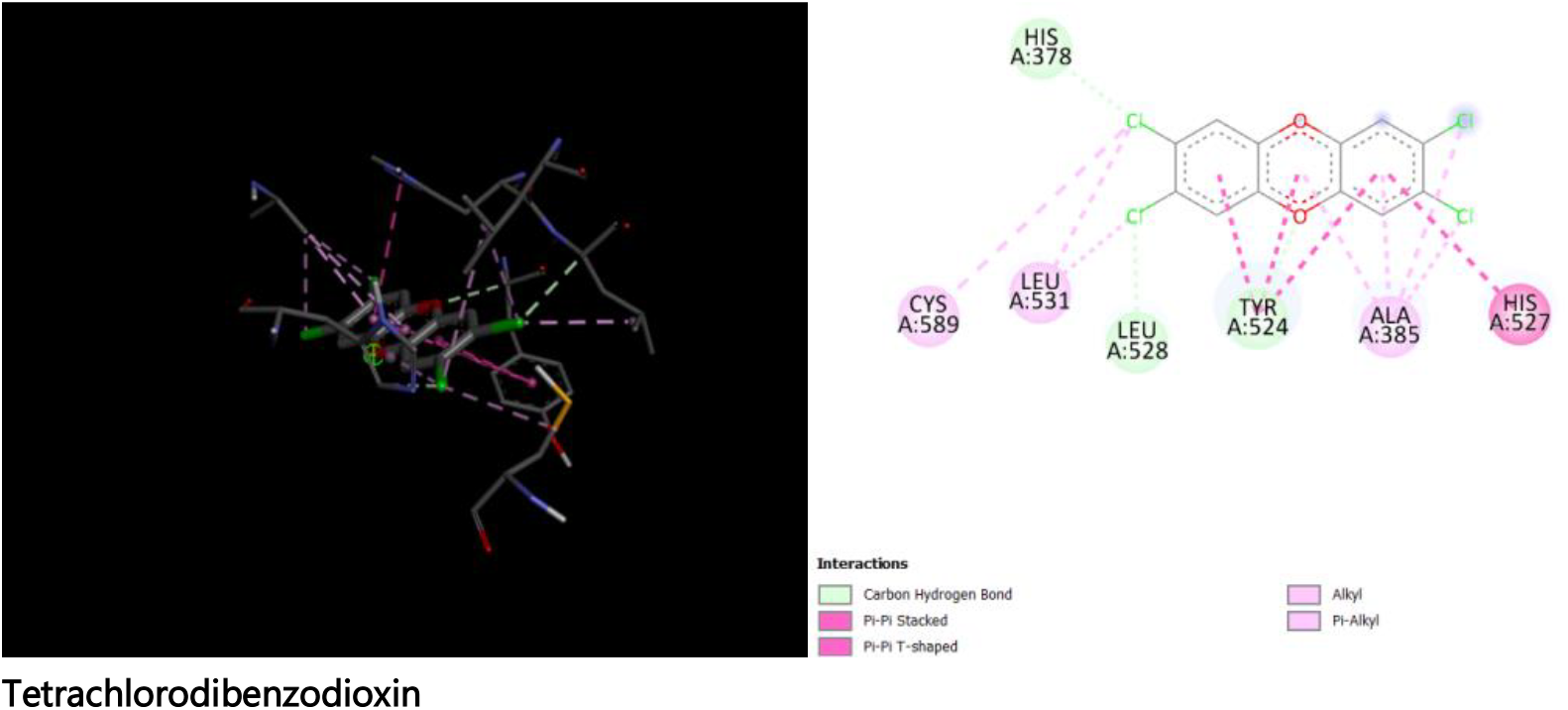
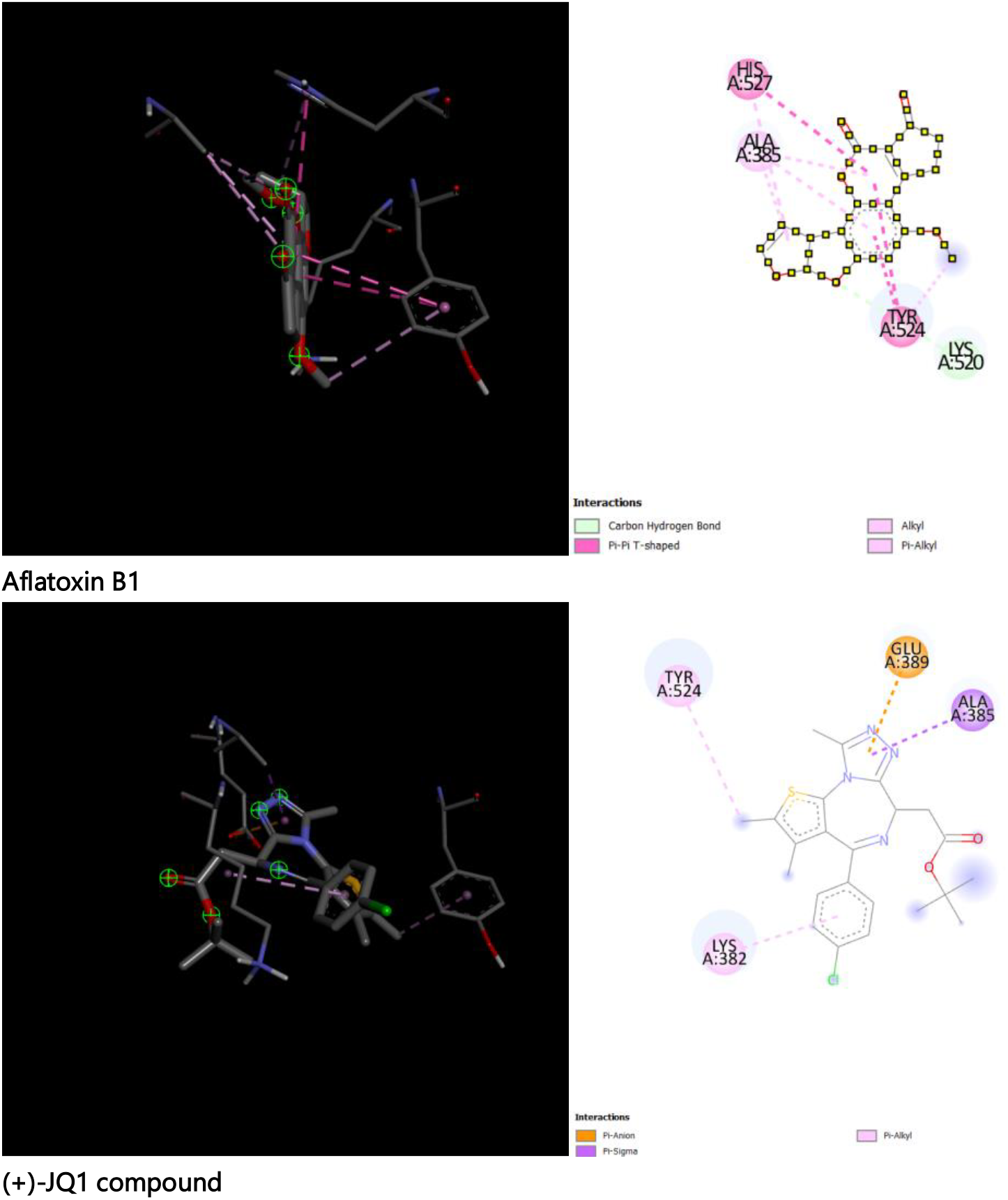
2D and 3D structure of Aflatoxin B1, (+)-JQ1 compound and Tetrachlorodibenzodioxin binding to *GBP1*

**Supplementary Figure 5.**
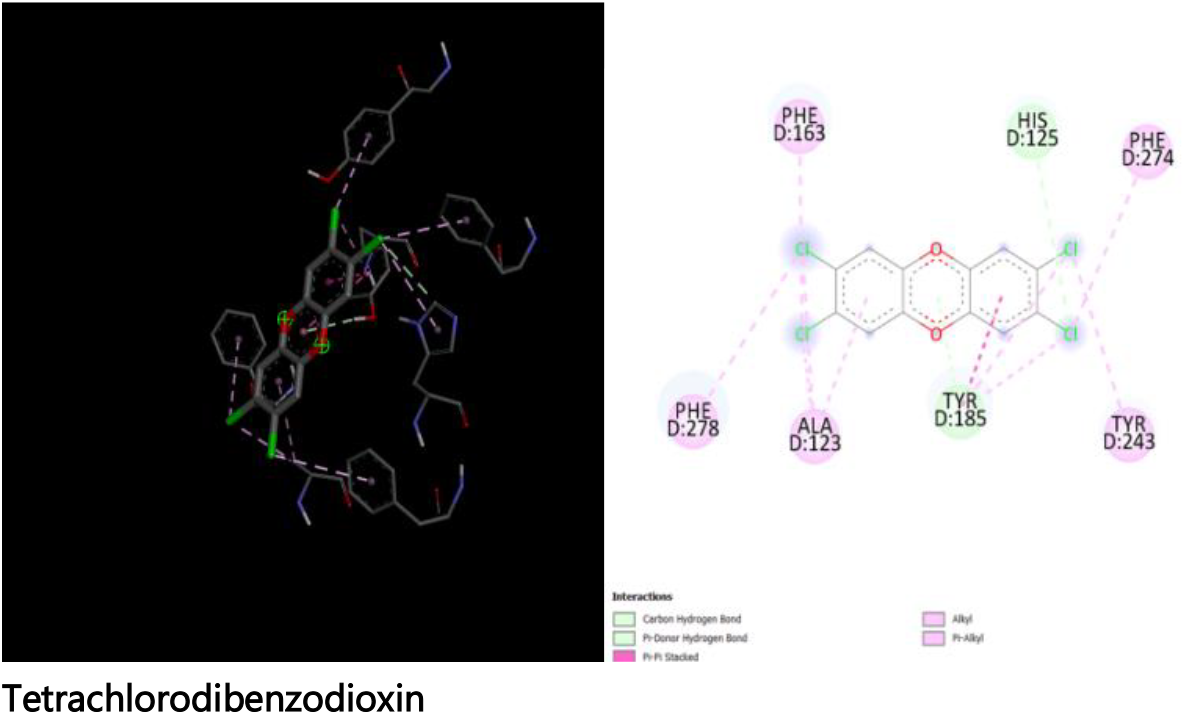
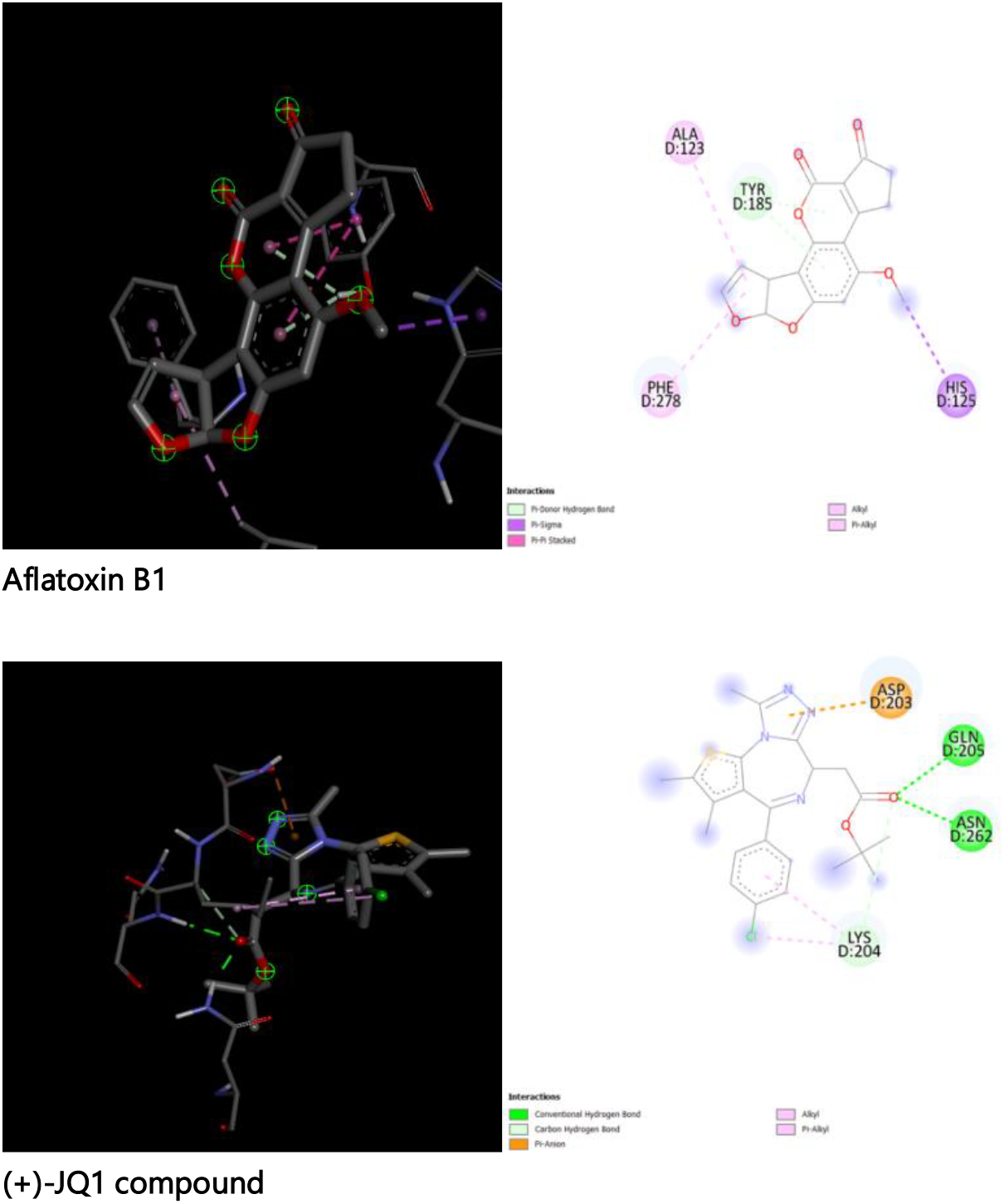
2D and 3D structure of Aflatoxin B1, (+)-JQ1 compound and Tetrachlorodibenzodioxin binding to TNFSF10

**Supplementary Figure 6.**
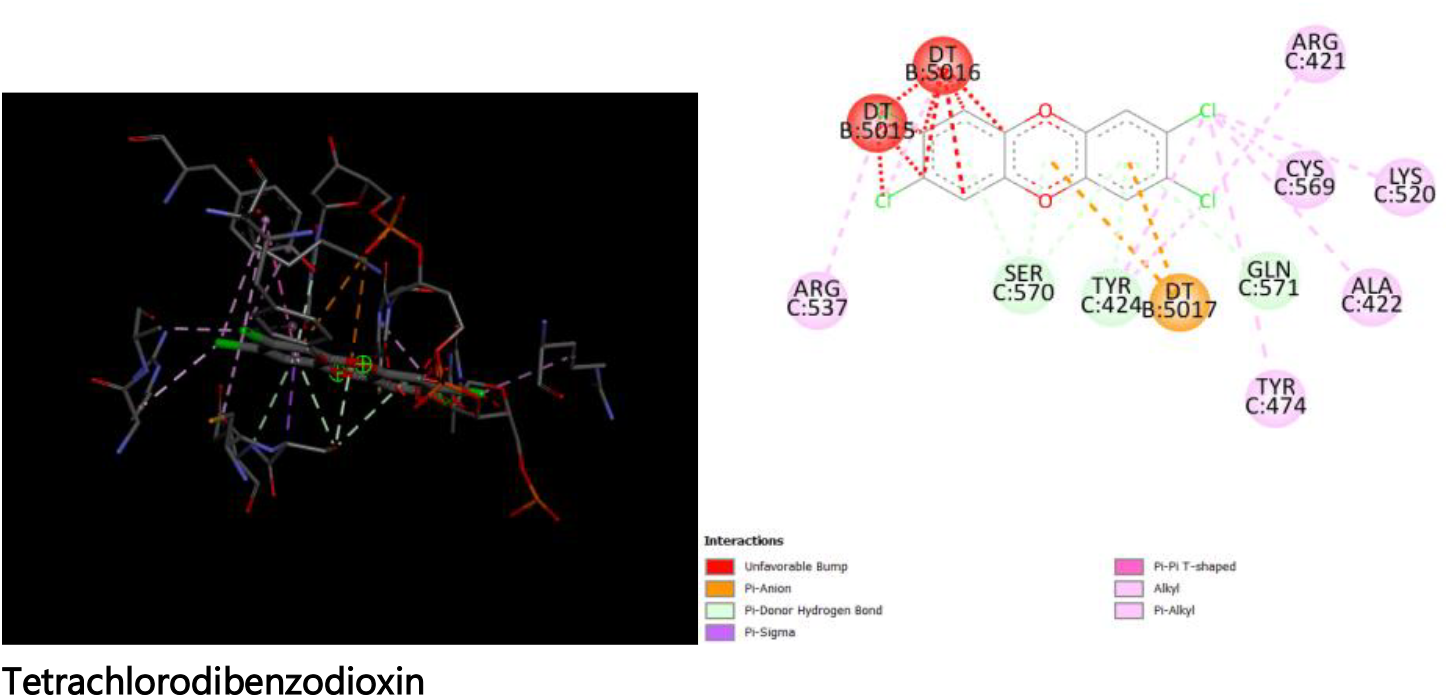
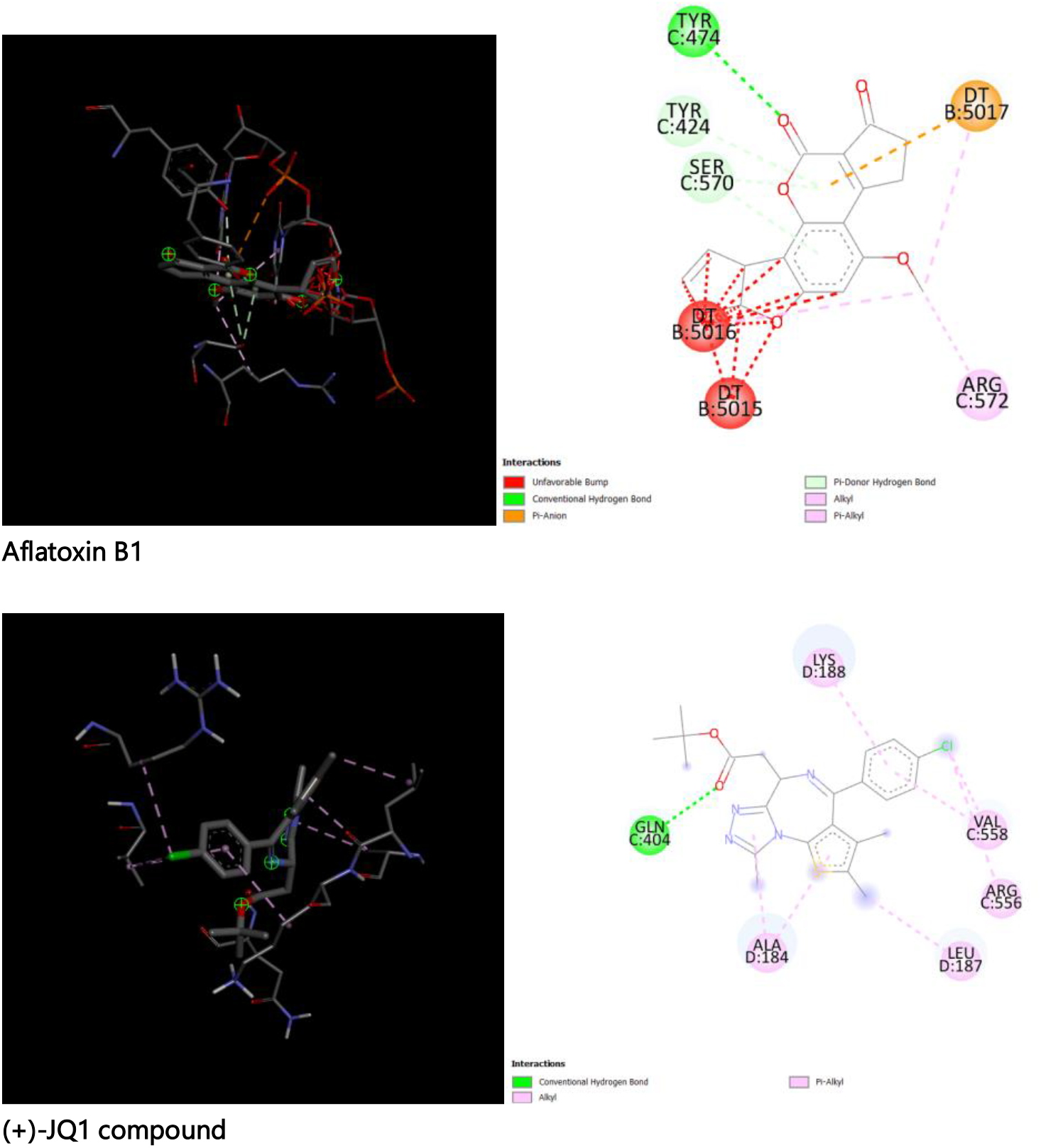
2D and 3D structure of Aflatoxin B1, (+)-JQ1 compound and Tetrachlorodibenzodioxin binding to *FOS*

**Supplementary Figure 7.**
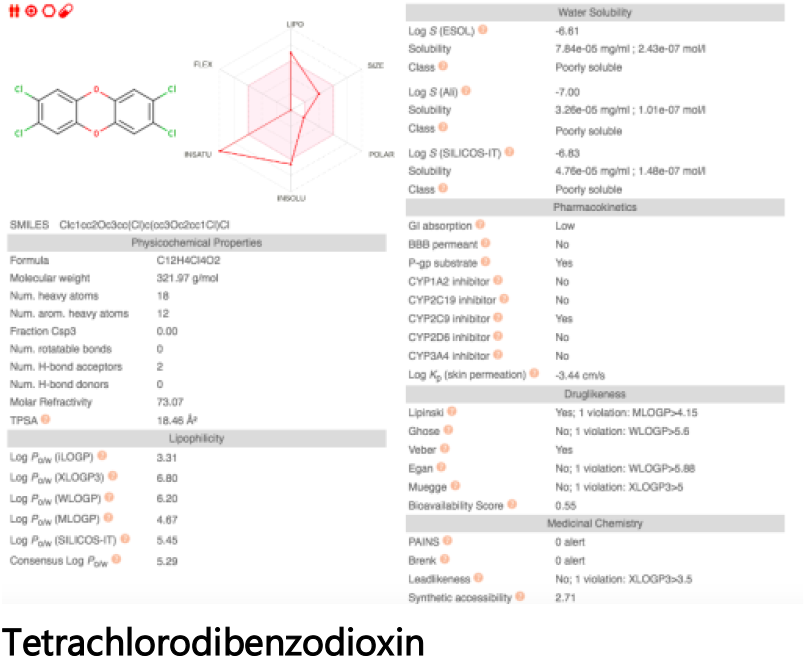
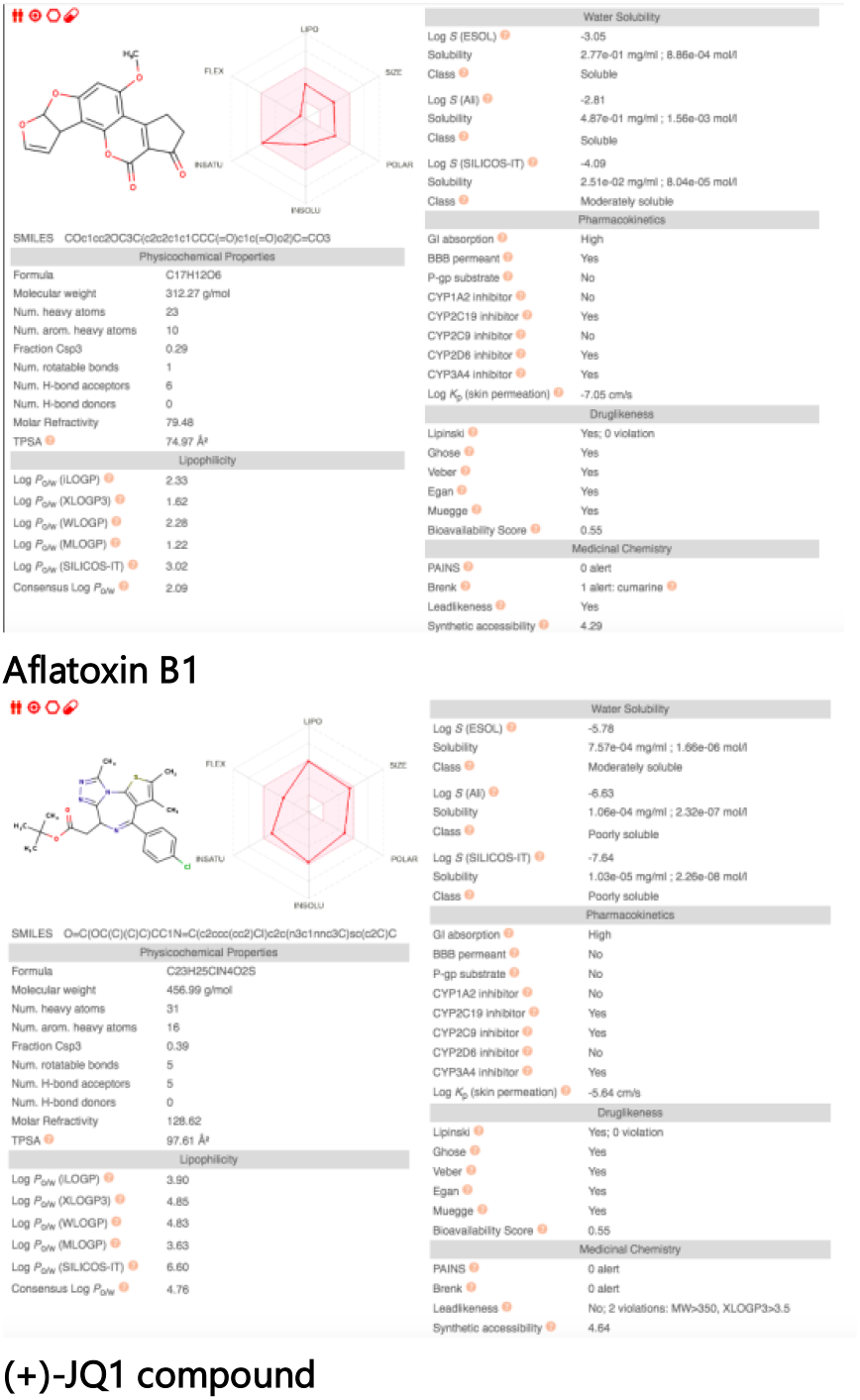
Druglikeness of Aflatoxin B1, (+)-JQ1 compound and Tetrachlorodibenzodioxin

